# The Lag3 KIEELE motif couples HPK1 to metabolic and functional regulation of Foxp3+ regulatory T cells

**DOI:** 10.64898/2026.09.28.755064

**Authors:** Giha Kim, Dongkyun Kim, Sohee Kim, Booki Min

## Abstract

Lymphocyte activation gene-3 (Lag3) is an inhibitory receptor that limits effector T cell activation and function. We previously demonstrated that Lag3 supports regulatory T (Treg) cell function through Myc-dependent metabolic reprogramming. However, the intracellular mechanisms underlying this regulation remain incompletely understood. Here, we generated mice with Treg-specific deletion of the Lag3 KIEELE motif and demonstrated that this motif is required for Treg-mediated suppression of autoimmune inflammation. Loss of the Lag3 KIEELE motif led to enhanced TCR/PI3K-Akt/MEK-ERK signaling and Myc expression, accompanied by increased glycolytic activity and acquisition of effector T cell-like features. We also identified that hematopoietic progenitor kinase 1 (HPK1) and Cbl-interacting protein of 85 kDa (CIN85) associate with Lag3 through the KIEELE motif. HPK1 depletion partially recapitulated the phenotypic changes and impaired Treg suppressive function in vivo. Together, these findings identify the KIEELE motif as a critical link between Lag3-associated signaling machinery and metabolic regulation that sustains Treg suppressive function during autoimmune inflammation.

## INTRODUCTION

Foxp3^+^ regulatory T (Treg) cells are essential for maintaining peripheral tolerance and limiting autoimmune inflammation (Sakaguchi et al., 2008). Their suppressive activity must be tightly regulated to prevent excessive immune activation while preserving protective immunity. Treg cells express a broad repertoire of immunoregulatory receptors, including PD-1, CTLA-4, ICOS, and Lag3, many of which are best characterized for their ability to limit conventional T cell activation and effector function (Chen and Flies, 2013; Joller et al., 2024). Increasing evidence, however, indicates that several of these receptors also play crucial roles in sustaining Treg cell fitness, stability, and suppressive function (Lucca et al., 2019; Wing et al., 2008; Francisco et al., 2009; Kim et al., 2024). Despite their importance in Treg cell biology, the precise mechanisms through which the receptors control Treg cell function remain poorly understood.

Lag3 has emerged as an important immune checkpoint receptor that restrains conventional T cell responses and a prominent therapeutic target for enhancing anti-tumor immunity, particularly in combination with blockade of the PD-1 pathway (Grosso et al., 2007; Woo et al., 2012; Tawbi et al., 2022). Beyond its well-established function in conventional T cells, Lag3 is expressed at high levels by Treg cells, although the mechanisms by which it regulates Treg cell function remain elusive. We previously reported that Lag3 expressed on Treg cells is required for effective control of autoimmune inflammation and that Lag3 restrains Myc-dependent metabolic reprogramming in Treg cells (Kim et al., 2024). These findings suggest that Lag3 supports Treg suppressive function, in part, by maintaining metabolic programs conducive to their optimal activity. However, the intracellular mechanisms linking Lag3 to the signaling pathways that control Treg cell metabolism and suppressive activity remain unknown. Elucidating the molecular machinery that couples Lag3 to intracellular signaling is therefore critical for understanding how it coordinates Treg cell metabolism and function to maintain immune homeostasis.

Unlike many co-inhibitory receptors, the cytoplasmic domain of Lag3 lacks canonical signaling motifs, such as immunoreceptor tyrosine-based inhibitory motifs (ITIMs) or immunoreceptor tyrosine-based switch motifs (ITSMs), suggesting that Lag3 engages distinct mechanisms to transduce inhibitory signals. Instead, the Lag3 cytoplasmic tail contains three conserved regions: an FxxL motif, a KIEELE motif, and a C-terminal glutamic acid-proline (EP) repeats (Chocarro et al., 2021; Maruhashi et al., 2020; Aggarwal et al., 2023). Previous studies examining the contribution of these motifs have yielded different results, primarily in conventional T cells. The FxxL motif has been implicated in Lag3-mediated inhibition independently of KIEELE (Maeda et al., 2019), whereas studies by Workman et al. identified the KIEELE motif as essential for Lag3-dependent suppression of T cell activation (Workman et al., 2002; Workman and Vignali, 2003). Lag3-associated protein (LAP) was initially reported to interact with the EP repeats (Iouzalen et al., 2001), but a functional requirement for LAP in Lag3-mediated signaling has not been established. More recently, Jiang et al. reported that the Cbl family proteins ubiquitinate a lysine residue within the KIEELE motif, promoting dissociation of the cytoplasmic tail from the plasma membrane and facilitating its inhibitory function in CD8^+^ T cells (Jiang et al., 2025). Nevertheless, how the KIEELE motif transduces signals that regulate cellular function and how such signaling is coupled to the metabolic programs that support Treg cell function are unknown.

Here, we investigated the role of the KIEELE motif in Lag3-dependent regulation of Treg function using a novel Treg cell-specific KIEELE-deficient mouse model. We found that the KIEELE motif is required for Treg cell-mediated control of autoimmune inflammation in vivo. Mechanistically, we identified hematopoietic progenitor kinase 1 (HPK1) and Cbl-interacting protein of 85 kDa (CIN85) as cytosolic proteins associated with the KIEELE motif. Loss of the KIEELE motif enhanced proximal TCR signaling and downstream PI3K-Akt and MEK-ERK pathways, driving metabolic reprogramming and IFNγ-dependent T-bet expression. Moreover, HPK1 knockdown in Treg cells recapitulated key metabolic and functional defects observed in KIEELE-deficient Treg cells. Together, these findings define the KIEELE motif as a key signaling component of Lag3 function in Treg cells and reveal a mechanism through which Lag3 restrains intracellular signaling and metabolic reprogramming to preserve Treg cell suppressive function.

## RESULTS AND DISCUSSION

### Generation of Treg cell-specific Lag3_KIEELE_^−/−^ mice

To investigate the role of the KIEELE motif in Lag3-dependent regulation of Treg cell function, we generated mice carrying a conditional Lag3 allele consisting of a floxed stop cassette followed by a mutant Lag3 sequence lacking the KIEELE motif (**Figure S1A**). These mice were crossed with Foxp3^CreYFP^ mice to selectively express the KIEELE-deficient Lag3 mutant in Treg cells. Genomic DNA sequencing of the Foxp3^CreYFP^ Lag3_KIEELE_^fl/fl^ (Treg^Lag3ΔKIEELE^) mice confirmed deletion of the KIEELE motif specifically in Tregs (**Figure S1B**). Treg^Lag3ΔKIEELE^ mice developed normally with total cellularity of CD4 and CD8 T cells in the secondary lymphoid tissues comparable to WT controls (**Figure S1C**). The frequency of CD44^high^ memory phenotype CD4 and CD8 T cells was also comparable (**Figure S1C**). Likewise, the frequency, total numbers, Foxp3 expression, and proliferation of CD4^+^ Foxp3^+^ Treg cells from Treg^Lag3ΔKIEELE^ mice were comparable to those of WT Treg cells (**Figure S1D and S1E**). Similarly, the expression of Treg cell-associated markers, ICOS, Nrp1, and GITR, was also comparable to WT Treg cells (**Figure S1F**). Together, these results indicate that the loss of the KIEELE motif of Lag3 does not impact Treg cell development or phenotype under steady-state condition.

### The KIEELE motif of Lag3 is critical for Treg cell suppressive function in vivo

To test the role of the KIEELE motif in Treg cell function in vivo, EAE was induced in Foxp3^CreYFP^ Lag3^WT/WT^ (Treg^WT^) and Treg^Lag3ΔKIEELE^ mice. Treg^Lag3ΔKIEELE^ mice developed more severe disease compared to Treg^WT^ mice (**Figure 1A**). This was represented by an increase in the frequency and number of CD4^+^ T cells infiltrating the CNS of Treg^Lag3ΔKIEELE^ mice compared to that of Treg^WT^ mice. Although the proportion of CNS-infiltrating Treg cells was comparable between the two groups, the absolute number was significantly higher in Treg^Lag3ΔKIEELE^ mice, consistent with the overall increase in CD4^+^ T cell accumulation (**Figure 1B**). Treg cells exhibited comparable expression of CD44, ICOS, GITR, CD25, Foxp3, and surface Lag3 across the two groups (**Figure S2A)**. CNS-infiltrating CD4^+^ T cells expressing pro-inflammatory cytokines, namely IL-17, TNFα, IFNγ, and GM-CSF, were significantly elevated in Treg^Lag3ΔKIEELE^ mice compared to Treg^WT^ mice (**Figure 1C**). Consistent with this finding, the mRNA expression of pro-inflammatory cytokines and chemokines was markedly increased in the CNS of Treg^Lag3ΔKIEELE^ mice (**Figure 1D**). To further characterize the Treg cell-intrinsic role of the KIEELE motif in Treg cell function, we performed an adoptive Treg cell transfer experiment where WT or CD4^cre^ Lag3_KIEELE_^fl/fl^ (Lag3_KIEELE_^−/−^) iTreg cells were transferred into EAE-induced mice depleted of endogenous Treg cells (**Figure S2B**). The recipients of Lag3_KIEELE_^−/−^ iTreg cells developed more severe disease and showed elevated number of CNS-infiltrating CD4^+^ T cells expressing inflammatory cytokines (**Figure 1E and 1F**). Collectively, these results demonstrate that the KIEELE motif is indispensable for Lag3-dependent Treg cell function to attenuate autoimmune inflammation.

**Figure 1.**
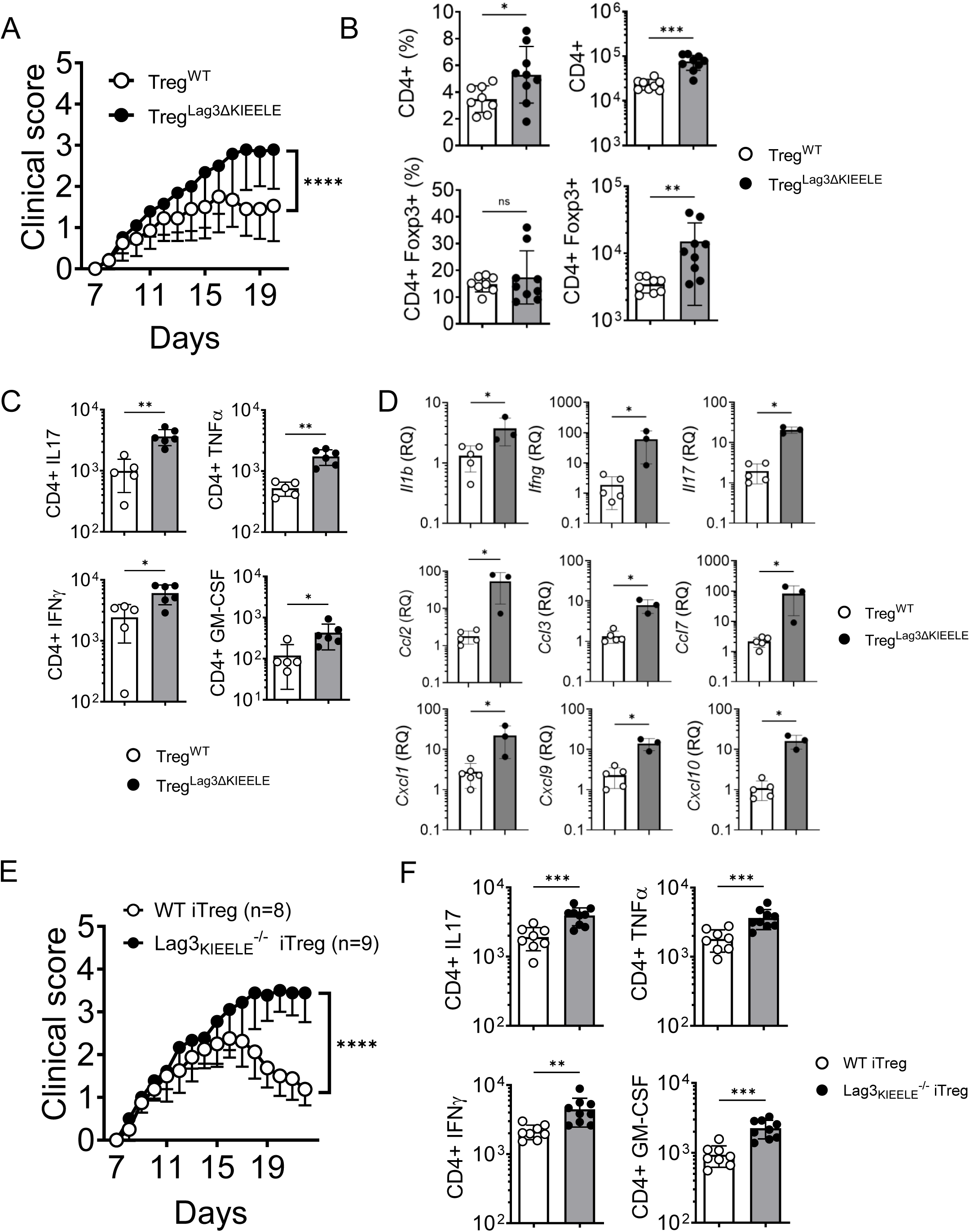
The Lag3 KIEELE motif is required for Treg-mediated suppression of autoimmune inflammation. (A) Clinical scores of Treg^WT^ and Treg^Lag3ΔKIEELE^ mice following EAE. (B) Frequency and total number of CD4^+^ T cells and CD4^+^ Foxp3^+^ Treg cells infiltrating the CNS at the peak of disease (day 14 post immunization). (C) Total number of IL-17-, TNFα-, IFNγ-, and GM-CSF-expressing CD4^+^ T cells infiltrating the CNS at the peak of disease. (D) mRNA expression measured from whole CNS tissue of mice induced with EAE. (E) Foxp3^DTR^ mice (n=8-9) were induced with EAE and treated with diphtheria toxin to deplete endogenous Tregs days 5 and 6 post immunization. Naïve CD4^+^ T cells from WT or Lag3_KIEELE_^−/−^ mice were polarized in Treg cell-polarizing condition for 3 days. 2x10^6^ WT or Lag3_KIEELE_^−/−^ iTregs cells were IV transferred day 7 post immunization. The mice were monitored and given clinical scores. (F) Total number of IL-17-TNFα-, IFNγ-, and GM-CSF-expressing CD4^+^ T cells infiltrating the CNS at the peak of disease. EAE clinical score data are presented as mean ± SEM. Data were pooled from four independent experiments in (A) (n = 19–20 mice per group) and two independent experiments in (E) (n = 8–9 mice per group). Statistical significance for (A) and (E) was assessed by two-way ANOVA, with genotype and day as factors. ****P < 0.0001 for the genotype × day interaction. Data from (B-D, F) were pooled from at least two independent experiments. Statistical significance for experiments in (B-D, F) was assessed using a two-tailed Mann–Whitney U test. *P < 0.05; **P < 0.01; ***P < 0.001; ****P < 0.0001.

### Loss of the Lag3 KIEELE motif enhances TCR signaling and drives metabolic reprogramming in Treg cells

We previously reported that Lag3 deficiency promotes Myc-dependent metabolic reprogramming that compromises Treg cell suppressive function. To determine whether KIEELE motif mediates similar metabolic regulation, we first measured Myc expression in WT and Lag3_KIEELE_^−/−^ iTreg cells. Indeed, Lag3_KIEELE_^−/−^ Treg cells exhibited increased expression of Myc and the glycolytic enzyme lactate dehydrogenase A (LDHA) at both the mRNA and protein levels compared with WT Treg cells (**Figure 2A and 2B**). This was further corroborated by an increase in extracellular acidification rate (ECAR) of Lag3_KIEELE_^−/−^ iTreg cells as measured by Seahorse analysis (**Figure 2C**). A similar increase in ECAR was observed in thymus-derived Lag3_KIEELE_^−/−^ tTreg cells (**Figure S3A**), indicating that KIEELE-dependent control of Treg cell metabolism is conserved in both induced and thymus-derived Treg cell populations.

**Figure 2.**
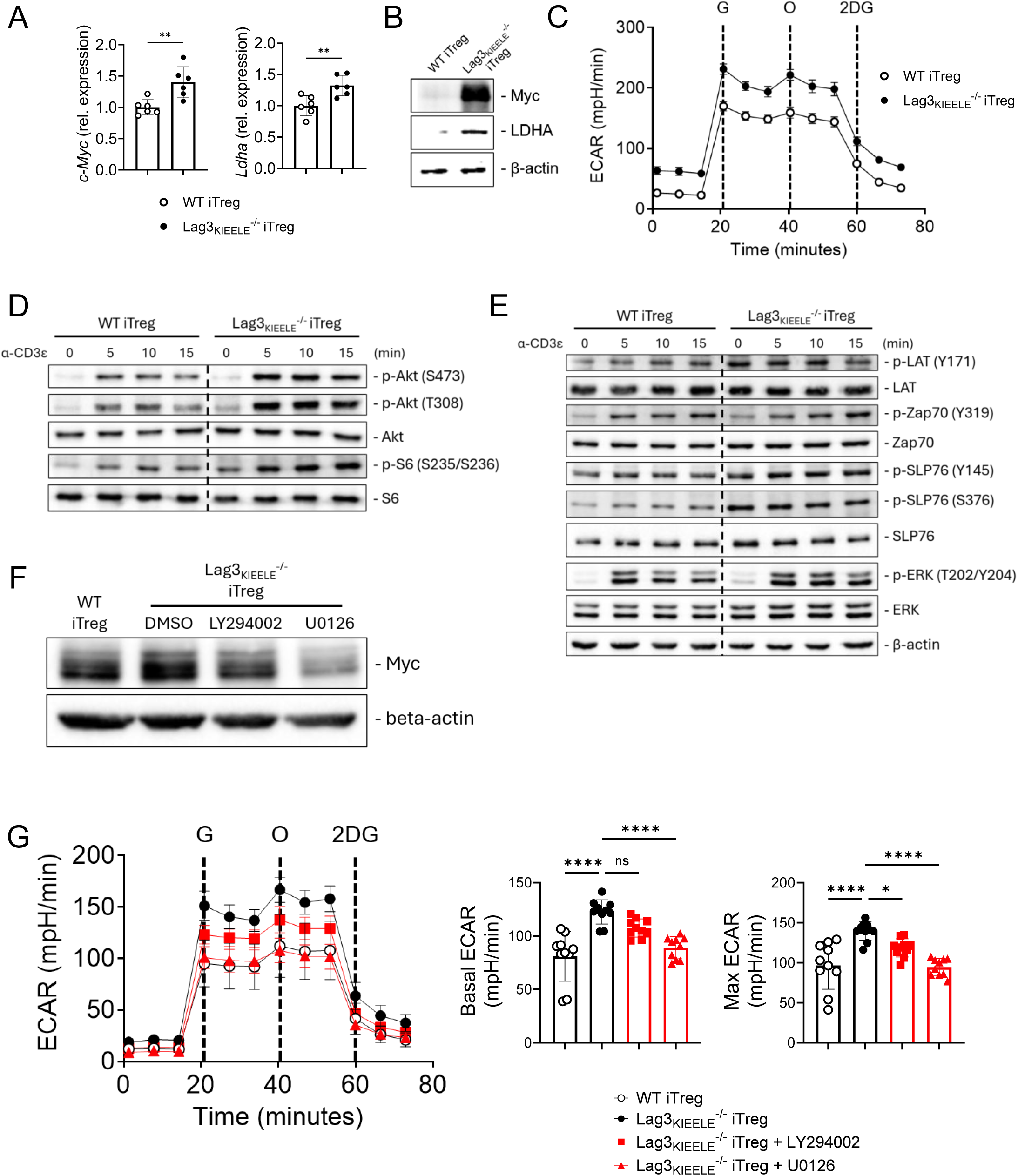
Deletion of the Lag3 KIEELE motif enhances TCR signaling and promotes metabolic reprogramming in Treg cells. (A) mRNA expression of *c-Myc* and *Ldha* measured from WT and Lag3_KIEELE_^−/−^ iTreg cells. (B) Myc and LDHA protein expression was assessed by western blotting. (C) ECAR of WT and Lag3_KIEELE_^−/−^ iTreg cells was measured by Seahorse. (D, E) WT or Lag3_KIEELE_^−/−^ iTreg cells were rested for 1 hr in serum-free RPMI and restimulated with biotinylated anti-CD3ε antibody cross-linked with streptavidin for the indicated times. Western blot analysis of p-Akt (S473 and T308), p-S6 (S235/236), total Akt and total S6 (D); and p-LAT (Y171), p-Zap70 (Y319), p-SLP76 (Y145 and S376), p-ERK (T202/Y204), and total LAT, Zap70, SLP76, and ERK (E). (F, G) Naïve CD4^+^ T cells from WT and Lag3_KIEELE_^−/−^ mice were polarized in Treg cell-polarizing condition for 3 days. Lag3_KIEELE_^−/−^ cells were treated with DMSO, LY294002, or U0126 during the final 24 h of culture. Myc protein expression was assessed by western blotting (F) and ECAR was measured by Seahorse (G). Data from (A-F) are representative of at least two independent experiments. Statistical significance for (A) was assessed using a two-tailed Mann–Whitney U test. Data from (G) were pooled from two independent experiments. Statistical significance was assessed by one-way ANOVA with Dunnett’s multiple-comparisons test. *P < 0.05; **P < 0.01; ****P < 0.0001; ns = not significant.

To examine the signaling mechanism underlying the metabolic reprogramming of Lag3_KIEELE_^−/−^ Treg cells, we examined pathways upstream of Myc. The PI3K-Akt axis is a major regulator of Myc expression and cellular metabolism with full Akt activation requiring phosphorylation at Thr308 and Ser473 and resulting in activation of downstream effectors, including S6. Following CD3ε stimulation, Lag3_KIEELE_^−/−^ iTreg cells exhibited increased phosphorylation of Akt at both residues, accompanied by enhanced S6 phosphorylation (**Figure 2D**), indicating elevated PI3K-Akt-mTOR pathway activation. Given the enhanced signaling, we next asked whether proximal TCR signaling was similarly altered in Lag3_KIEELE_^−/−^ Treg cells. Following CD3ε stimulation, Lag3_KIEELE_^−/−^ Treg cells exhibited increased phosphorylation of the proximal TCR signaling molecules, LAT and Zap70, as well as the downstream kinase ERK (**Figure 2E**). Phosphorylation of SLP76 was also elevated at both Tyr145 and Ser376 (**Figure 2E**). While Tyr145 phosphorylation promotes TCR signaling, phosphorylation at Ser376 has been associated with attenuation of TCR signaling (Jordan et al., 2006; Bartolo et al., 2007). HPK1 phosphorylates SLP76 at Ser376, promoting the recruitment of 14-3-3 proteins, and has also been implicated in SLP76 ubiquitination and proteasomal degradation, potentially through the recruitment of an unidentified E3 ubiquitin ligase (Wang et al., 2012). However, SLP76 Ser376 phosphorylation was increased in Lag3_KIEELE_^−/−^ iTregs, and this was not accompanied by a reduction in SLP76 protein abundance (**Figure S3B**). In contrast, Myc underwent similarly rapid degradation in both genotypes within the first 2 hours, consistent with previous reports (Farrell and Sears, 2014). Therefore, SLP76 Ser376 phosphorylation alone appears to be insufficient to promote SLP76 degradation in Treg cells, suggesting that additional regulatory mechanisms may determine the fate of phosphorylated SLP76. Next, we sought to determine whether enhanced PI3K-Akt and MEK-ERK signaling contributed to Myc upregulation and the associated metabolic phenotype. Pharmacological inhibition of either PI3K with LY294002 or MEK with U0126 markedly reduced Myc expression in Lag3_KIEELE_^−/−^ Treg cells, with MEK inhibition exerting a more pronounced effect (**Figure 2F**). Inhibition of either pathway reduced ECAR in Lag3_KIEELE_^−/−^ Treg cells, with greater reduction following MEK inhibition paralleling its stronger suppression of Myc expression (**Figure 2G**). Together, these results suggest that the KIEELE motif restrains TCR-dependent PI3K-Akt and MEK-ERK signaling, thereby limiting Myc induction and glycolytic reprogramming in Treg cells.

### Lag3 KIEELE motif recruits key proteins to restrain TCR-PI3K signaling and sustain Treg cell suppressive function

We next sought to define the molecular basis for the enhanced proximal TCR and PI3K signaling observed in Lag3_KIEELE_^−/−^ Treg cells. A recent study demonstrated that the lysine residue within the KIEELE motif is ubiquitinated by the E3 ubiquitin ligases, c-Cbl and Cbl-b, thereby disrupting the binding of the Lag3 cytoplasmic tail to the membrane and stabilizing it in a membrane-dissociated conformation (Jiang et al., 2025). This observation prompted us to investigate whether Cbl-associated proteins might link the KIEELE motif to the regulation of TCR and PI3K signaling. CIN85 emerged as a candidate because of its ability to associate with Cbl proteins and its reported involvement in the regulation of TCR signaling pathways (Kong et al., 2019; Luff et al., 2021). To determine whether CIN85 associates with the Lag3 cytoplasmic domain, we performed co-immunoprecipitation using lysates from WT, CD4^cre^ Lag3_CyD_^fl/fl^ (Lag3_CyD_^−/−^), or Lag3_KIEELE_^−/−^ iTreg cells. Following Lag3 immunoprecipitation, CIN85 was readily detected in WT Treg cells but was absent in Lag3_CyD_^−/−^ cells (**Figure 3A**), demonstrating that its association with Lag3 requires the cytoplasmic domain.

**Figure 3.**
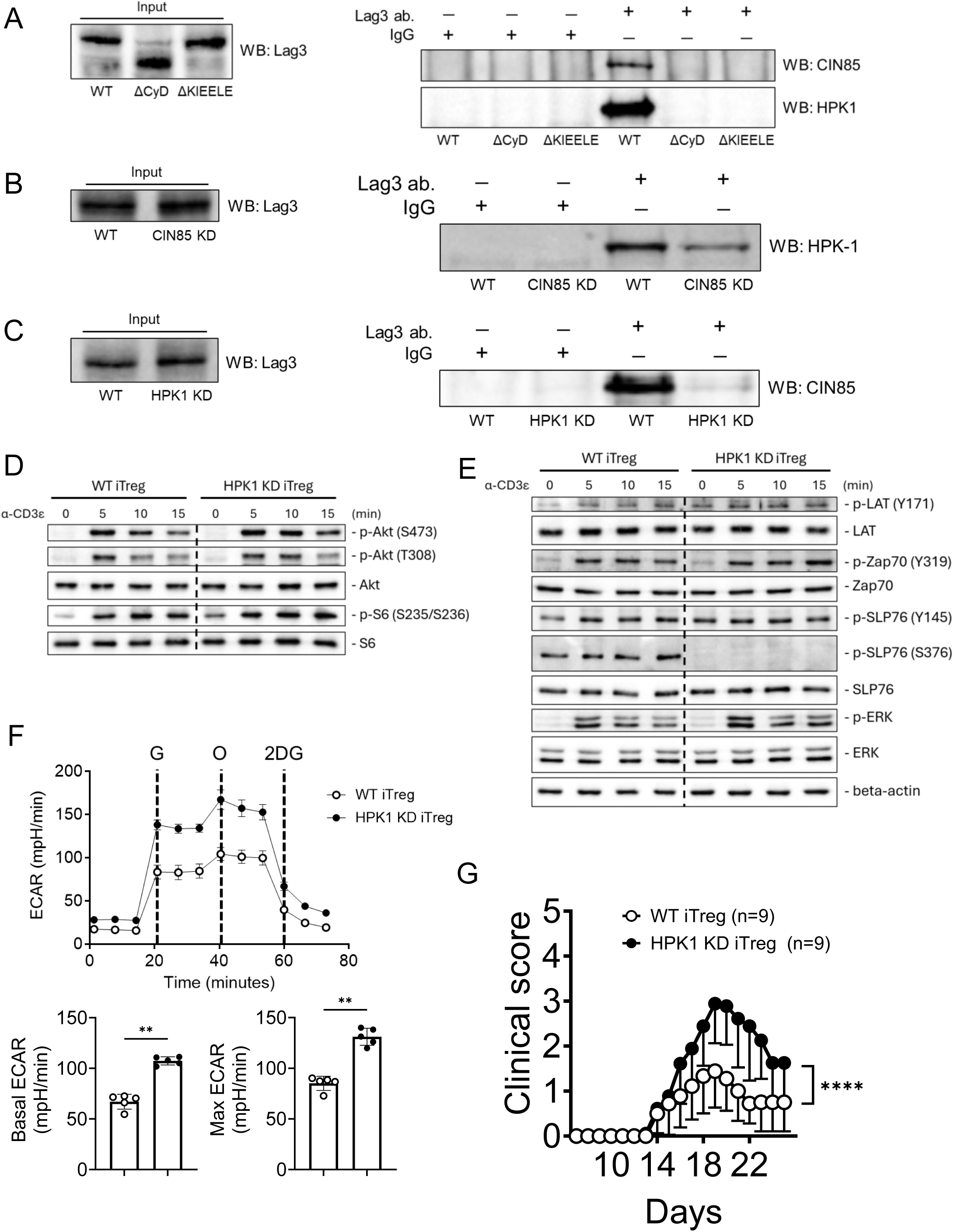
HPK1 and CIN85 associate with Lag3 in a KIEELE-dependent manner, and HPK1 depletion enhances TCR signaling and metabolic reprogramming while impairing Treg suppressive function. (A-C) Lysates from WT, Lag3_CyD_^−/−^, and Lag3_KIEELE_^−/−^ iTreg cells (A), WT and CIN85 KD iTreg cells (B), or WT and HPK1 KD iTreg cells (C) were immunoprecipitated with anti-Lag3 or an isotype-matched IgG control and analyzed by western blotting for CIN85 and HPK1. Input lysates were analyzed for Lag3. (D, E) WT or HPK1 KD iTreg cells were rested for 1 hr in serum-free RPMI and restimulated with biotinylated anti-CD3ε antibody cross-linked with streptavidin for the indicated times. Western blot analysis of p-Akt (S473 and T308), p-S6 (S235/236), total Akt and total S6 (D); and p-LAT (Y171), p-Zap70 (Y319), p-SLP76 (Y145 and S376), p-ERK (T202/Y204), and total LAT, Zap70, SLP76, and ERK (E). (F) ECAR of WT and HPK1 KD iTreg cells was measured by Seahorse. (G) Foxp3^DTR^ mice were induced with EAE and treated with diphtheria toxin to deplete endogenous Tregs days 5 and 6 post immunization. 2x10^6^ WT or HPK1 KD iTregs cells were IV transferred day 10 post immunization. Data from (A-F) are representative of at least two independent experiments. Data from (G) are presented as the mean clinical score ± SEM and are combined from two independent experiments (n = 9 mice per group). Statistical significance was determined by two-tailed Mann– Whitney U test in (F) and two-way repeated-measures ANOVA in (G), with Treg treatment group (WT or HPK1 KD) and day as factors. **P < 0.01; ****(P < 0.0001) for the treatment group × day interaction.

Importantly, CIN85 was also undetectable in Lag3_KIEELE_^−/−^ Treg cells, indicating that CIN85 associates with Lag3 through the KIEELE motif (**Figure 3A**). Having identified CIN85 as a KIEELE-dependent Lag3-associated protein, we next asked whether additional signaling proteins might participate in this complex. Among previously identified CIN85-interacting proteins, HPK1 was of particular interest because of its established function as a negative regulator of TCR signaling (Büchse et al., 2011; Shui et al., 2007; Hernandez et al., 2018). We thus tested whether HPK1 also associates with the Lag3 cytoplasmic domain. Indeed, HPK1 co-immunoprecipitated with Lag3 in WT Treg cells, whereas this association was lost in both Lag3_CyD_^−/−^ and Lag3_KIEELE_^−/−^ Treg cells (**Figure 3A**). We next tested whether the associations of CIN85 and HPK1 with Lag3 are interdependent using a sgRNA-based CRISPR-Cas9 knockdown approach. Naïve CD4 T cells were subjected to CRISPR-Cas9-mediated knockdown of CIN85 or HPK1 and subsequently differentiated into iTreg cells. Knockdown was validated by western blot analysis (**Figure S4A and S4B**). Moreover, the knockdown did not affect the generation of Treg cells (**Figure S4C and S4D**). HPK1 remained detectable in Lag3 immunoprecipitates following CIN85 knockdown, indicating that association of HPK1 with Lag3 does not require CIN85 (**Figure 3B**). In contrast, CIN85 was no longer detected in Lag3 immunoprecipitates following HPK1 knockdown (**Figure 3C**), indicating that the association of CIN85 with Lag3 is dependent on HPK1. Because both HPK1 and CIN85 have been implicated in the negative regulation of TCR and PI3K signaling, we hypothesized that disruption of this KIEELE-associated complex would contribute to the enhanced signaling observed in Lag3_KIEELE_^−/−^ Treg cells. Given that CIN85 association with Lag3 was dependent on HPK1, we focused on defining the functional contribution of HPK1. HPK1 KD iTreg cells exhibited enhanced PI3K-Akt and proximal TCR signaling compared with WT iTreg cells, resembling the signaling phenotype observed in Lag3_KIEELE_^−/−^ Treg cells, although the increases were less pronounced (**Figure 3D and 3E**). Consistent with efficient CRISPR-mediated knockdown, HPK1 depletion completely abolished SLP76 Ser376 phosphorylation (**Figure 3E**). However, despite the loss of Ser376 phosphorylation, there was no detectable increase in total SLP76 protein abundance, suggesting that phosphorylation at this site does not necessarily control SLP76 degradation in Treg cells (**Figure 3E**).

HPK1 KD Treg cells also displayed increased ECAR, further phenocopying the metabolic alterations associated with loss of the KIEELE motif (**Figure 3F**). Finally, we assessed whether HPK1 is required for Treg cell suppressive function in vivo. WT or HPK1 KD iTreg cells were adoptively transferred into Treg cell-depleted mice subjected to EAE. Whereas transfer of WT Treg cells effectively limited disease severity, mice receiving HPK1 KD Treg cells developed more severe EAE (**Figure 3G**), indicating that HPK1 is required for optimal Treg cell-mediated control of autoimmune inflammation.

Therefore, these findings suggest that the KIEELE motif enables the association of HPK1 with the Lag3 cytoplasmic domain, which in turn facilitates CIN85 recruitment. Disruption of this pathway enhances proximal TCR and PI3K-Akt signaling, promotes glycolytic reprogramming, and compromises Treg cell suppressive function.

### Lag3 regulates Teff cell-signature in Treg cells through the KIEELE motif

Treg cells exhibit substantial phenotypic plasticity in response to signals within the local microenvironment and can acquire effector T cell (Teff)-like features under inflammatory conditions (Koch et al., 2009; Dominguez-Villar et al., 2011; Feng et al., 2011; Voo et al., 2009; Kim et al., 2017). Given that Lag3_KIEELE_^−/−^ Treg cells displayed enhanced TCR and PI3K signaling together with increased Myc expression and glycolytic activity, features associated with Teff cells, we next tested whether loss of the KIEELE motif predisposes Treg cells to acquire a Teff-like phenotype. Indeed, Lag3_KIEELE_^−/−^ Treg cells exhibited increased expression of *Tbx21* and *Ifng* mRNA compared with WT Treg cells (**Figure 4A**). Likewise, flow cytometric analysis revealed increased T-bet expression in Lag3_KIEELE_^−/−^ Treg cells, which was further enhanced by IFNγ stimulation (**Figure 4B and S5A**). Inhibition of either PI3K with LY294002 or MEK with U0126 reduced T-bet expression, with MEK inhibition producing a more pronounced effect (**Figure 4B and S5A**), suggesting that heightened intracellular signaling downstream of Lag3 may contribute to the acquisition of this phenotype. Given our identification of HPK1 as a KIEELE-dependent Lag3-associated protein that restrains TCR signaling, we examined whether HPK1 knockdown similarly affected T-bet expression. HPK1 KD iTreg cells exhibited increased *Tbx21* and *Ifng* mRNA and T-bet protein expression, consistent with contribution of HPK1 to Lag3-mediated restraint of T-bet expression (**Figure S5B and S5C**).

**Figure 4.**
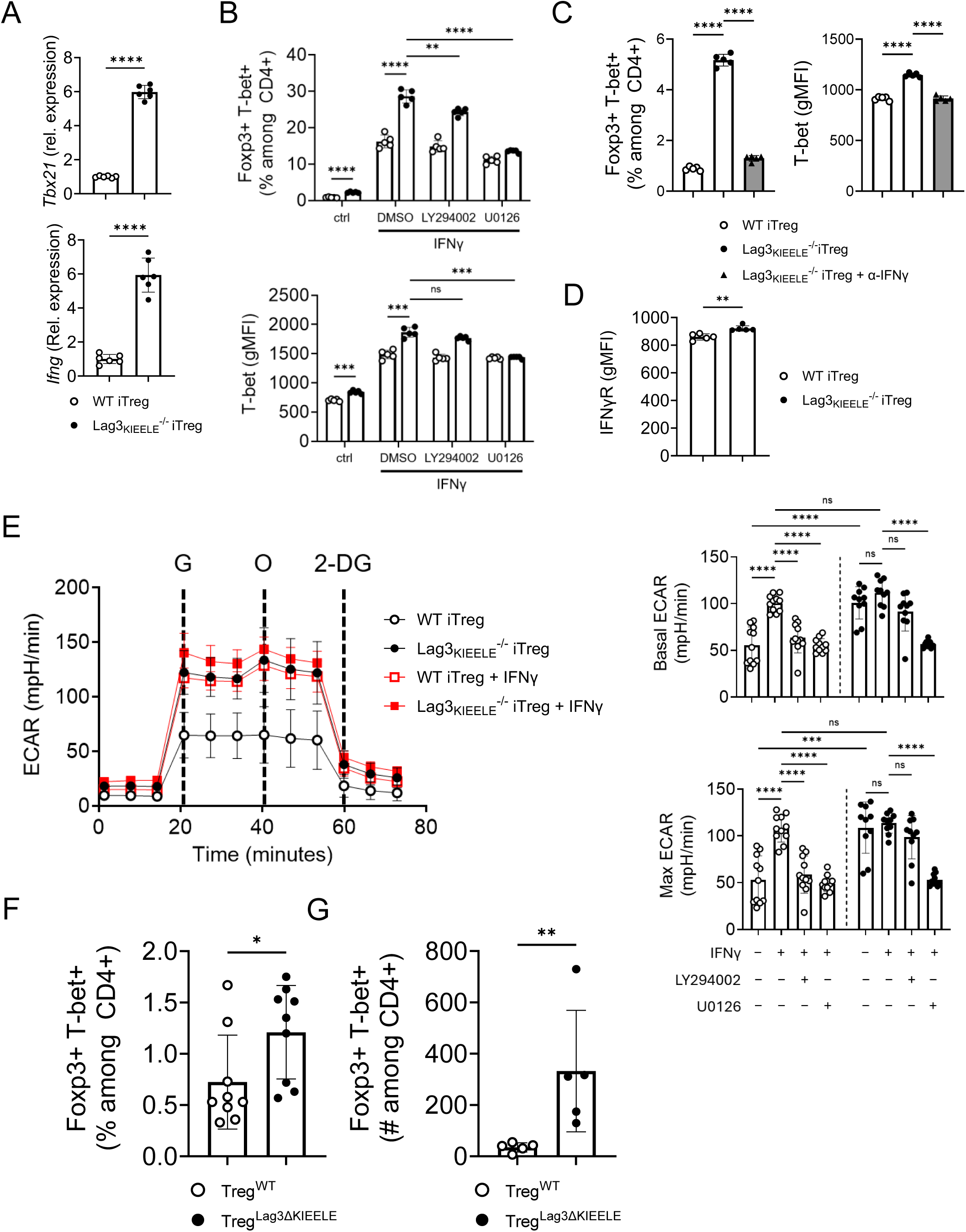
KIEELE-deficient Treg cells exhibit increased T-bet expression and altered responses to IFNγ. (A) mRNA expression of *Tbx21* and *Ifng* measured from WT and Lag3_KIEELE_^−/−^ iTreg cells. (B, C, E) Naïve CD4^+^ T cells from WT and Lag3_KIEELE_^−/−^ mice were cultured under Treg-polarizing conditions for 3 days. For (B, E), cells were treated with IFNγ in the presence or absence of DMSO, LY294002, or U0126 during the final 24 h of culture. T-bet expression was assessed by flow cytometry (B) and ECAR was measured by Seahorse (E). For (C), cells were cultured with or without IFNγ-neutralizing antibodies throughout the 3-day polarization period, and T-bet expression was assessed by flow cytometry. (D) IFNγR expression on WT and Lag3_KIEELE_^−/−^ iTreg cells was assessed by flow cytometry. (F, G) EAE was induced in Treg^WT^ and Treg^Lag3ΔKIEELE^ mice. T-bet expression in CNS-infiltrating Treg cells (F) and absolute numbers of CNS-infiltrating Foxp3^+^ T-bet^+^ cells (G) were determined by flow cytometry. Data from (A-D) are representative of at least two independent experiments and data from (E-G) are pooled from two independent experiments. Statistical significance was determined by two-tailed unpaired t test with Welch’s correction in (A) and (D), two-way repeated-measures ANOVA with Šídák’s multiple comparisons test in (B) and (E), one-way ANOVA with Dunnett’s multiple-comparisons test in (C), and two-tailed Mann–Whitney U test in (F) and (G). *P < 0.05; **P < 0.01; ***P < 0.001; ****P < 0.0001; ns = not significant.

We next investigated the contribution of IFNγ to T-bet expression. Although IFNγ increased T-bet expression in both WT and Lag3_KIEELE_^−/−^ iTreg cells, the magnitude of T-bet induction was substantially greater in Lag3_KIEELE_^−/−^ cells (**Figure 4B**). Since these cells also express increased *Ifng* expression, we hypothesized that Treg cell-derived IFNγ may reinforce T-bet expression through an autocrine positive-feedback loop.

Indeed, neutralization of IFNγ markedly reduced T-bet expression in Lag3_KIEELE_^−/−^ Treg cells (**Figure 4C and S5D**). Lag3_KIEELE_^−/−^ Treg cells expressed modestly but significantly higher levels of IFNγR1 than WT Treg cells (**Figure 4D**), potentially enhancing their responsiveness to IFNγ. We examined the effect of IFNγ on Treg cell metabolism. IFNγ treatment markedly increased ECAR in WT Treg cells to levels comparable to those observed in Lag3_KIEELE_^−/−^ Treg cells (**Figure 4E**), suggesting that IFNγ promotes glycolytic programming in Treg cells. In contrast, IFNγ did not further increase ECAR in Lag3_KIEELE_^−/−^ iTregs, potentially reflecting their already elevated glycolytic state.

Importantly, the IFNγ-induced increase in ECAR in WT Treg cells was reversed by either LY294002 or U0126 (**Figure 4E**), indicating that PI3K and MEK-ERK signaling may contribute to IFNγ-mediated metabolic reprogramming. Lastly, we determined whether the Teff-like phenotype observed in vitro was also found under inflammatory conditions in vivo. Following EAE induction, CNS-infiltrating Treg cells were isolated from Treg^WT^ or Treg^Lag3ΔKIEELE^ mice and T-bet expression was assessed by flow cytometry. Consistent with our in vitro findings, both the frequency and absolute number of T-bet^+^ CNS-infiltrating Treg cells were increased in Treg^Lag3ΔKIEELE^ mice compared with WT controls (**Figure 4F, 4G and S5E**). Collectively, these results suggest that the Lag3 KIEELE motif restrains the acquisition of Teff-like features in Treg cells by limiting proximal PI3K and MEK-ERK signaling and an IFNγ-dependent program associated with increased T-bet expression and glycolytic reprogramming.

Our findings identify a novel mechanism by which the Lag3 cytoplasmic domain controls Treg cell metabolism and function. Specifically, the KIEELE motif promotes the association of Lag3 with HPK1 and CIN85, providing a molecular link between Lag3 and the proximal TCR and downstream PI3K-Akt and MEK-ERK pathways. Our coimmunoprecipitation data further suggest a model in which HPK1 functions as a scaffold or adaptor that facilitates the recruitment of CIN85 and potentially other regulatory proteins to the Lag3 cytoplasmic domain. Given the established roles of HPK1 and CIN85 in negatively regulating TCR signaling, these interactions likely provide a mechanism by which the KIEELE motif restrains excessive TCR-associated signaling in Treg cells. Although our data suggest that HPK1 is necessary for the CIN85 and KIEELE motif interaction, whether HPK1 directly interacts with the KIEELE motif, or alternatively, recruits additional components that organize the Lag3-associated complexes will require further investigation. Whether similar or distinct inhibitory signaling pathway operates in different cells, especially conventional T cells, is under investigation.

Treg cells adapt to inflammatory and tissue-specific environments by acquiring distinct transcriptional programs, including expression of transcription factors associated with effector T cell lineages (Campbell and Koch, 2011; Panduro et al., 2016; Malviya et al., 2023). T-bet expression can promote Treg cell adaptation to facilitate the control of Th1-type responses (Koch et al., 2009; Dominguez-Villar et al., 2011). Our results demonstrate that T-bet can be induced through the combination of increased TCR, PI3K-Akt, and MEK-ERK signaling, together with enhanced glycolysis and an IFNγ-dependent mechanism. Our results also suggest that T-bet expression alone does not define the functional state of Treg cells; rather, its effects may depend on the specific signaling and metabolic pathways active in the cells. Therefore, in Lag3_KIEELE_^−/−^ Treg cells, this coordinated shift toward enhanced signaling, glycolytic metabolism, and IFN-γ responsiveness may promote a Teff-like state that is less compatible with optimal suppressive function. These findings highlight the context-dependent nature of Treg cell plasticity and suggest that Lag3 signaling helps constrain effector-associated programs that can destabilize Treg cell function.

In summary, our findings identify HPK1 and CIN85 as KIEELE-dependent Lag3-associated proteins that contribute to the restraint of excessive signaling in Treg cells. Disruption of KIEELE-dependent regulation enhanced proximal TCR, PI3K-Akt, and MEK-ERK signaling, leading to metabolic reprogramming and IFNγ-dependent T-bet expression and ultimately compromising their suppressive function. Together, our study provides a mechanistic framework for Lag3-mediated intracellular signaling and establishes the KIEELE motif as a critical link between Lag3, intracellular signaling, metabolic fitness, and suppressive function in Treg cells.

## MATERIALS AND METHODS

### Mice

Foxp3CreYFP (B6.129(Cg)-Foxp3tm4(YFP/icre)Ayr/J, #016959) and Foxp3DTR (B6.129(Cg)-Foxp3tm3(DTR/GFP)Ayr/J, #016958) were purchased from the Jackson Laboratory (Bar Harbor, ME). Lag3_KIEELE_-floxed mice were generated by Biocytogen (Beijing, China). Both male and female mice were used between 6 and 12 weeks of age. All the mouse protocols were approved by the Institutional Animal Care and Use Committee (IACUC) of Northwestern University. The mice were maintained under specific pathogen-free conditions at the Northwestern University Feinberg School of Medicine.

### EAE induction

Mice were immunized subcutaneously in the rear flanks with 1:1 emulsion of MOG_35-55_ peptide (300 μg) and complete Freund’s adjuvant (CFA) containing 5 mg/ml of Mycobacterium tuberculosis H37Ra. Pertussis toxin (200 ng) was administered intraperitoneally on days 0 and 2 after immunization. For adoptive transfer, Foxp3-DTR mice received diphtheria toxin intraperitoneally on days 5 and 6 post-immunization, followed by intravenous (IV) transfer of 2x10^6^ Treg cells of the indicated genotype or treatment groups. EAE severity was assessed using the following clinical scoring scale: 0 = No clinical signs; 1 = Limp tail; 2 = Paraparesis (weakness, incomplete paralysis of one or two hind limbs); 3 = Paraplegia (complete paralysis of two hind limbs); 4 = Paraplegia with forelimb weakness or paralysis; 5 = Moribund state or death.

### Isolation and culture of primary T cells

Primary CD4^+^ T cells were isolated from the spleens and lymph nodes of mice of the indicated genotypes. Tissues were mechanically dissociated into single-cell suspensions and passed through a 70-μm cell strainer. Splenic erythrocytes were lysed using Ack lysis buffer. Naïve CD4^+^ CD44^lo^ CD25^−^ cells were sorted and stimulated with plate-bound anti-CD3ε (2C11) and anti-CD28 (37.51) and cultured in the presence of recombinant TGF-β1 (5 ng/ml) and IL-2 (100 U/ml) for 3 days. Where indicated, cells were treated with recombinant IFNγ (50 ng/ml), LY294002 (5 µM), U0126 (5 µM), or DMSO vehicle during the final 24 h of culture. For IFNγ neutralization, anti-IFNγ antibody (XMG1.2; 10 µg/ml) was added at the initiation of culture and maintained throughout the 3-day polarization period.

### TCR stimulation

iTreg cells (5x10^5^) of the indicated genotypes were rested in serum-free RPMI for 1 h at 37°C. Cells were then pelleted by centrifugation, resuspended in serum-free RPMI, and maintained on ice while biotinylated anti-CD3ε antibody (5 µg/ml) and streptavidin (20 µg/ml) were added sequentially. TCR stimulation was initiated by transferring cells to 37°C for the indicated durations. Stimulation was terminated by immediate addition of Laemmli sample buffer and lysates were analyzed by western blotting.

### Flow cytometry

Single-cell suspensions were obtained from in vitro cultures or prepared from brain and spinal cord tissues as previously described (Kim et al., 2024). Cells were stained with fluorochrome-conjugated antibodies against surface markers for 30 min at 4°C in the dark. Dead cells were excluded using Zombie Violet Fixable Viability Kit (BioLegend) applied according to the manufacturer’s instructions. For intracellular staining, cells were fixed and permeabilized using Foxp3/transcription factor staining buffer set (Invitrogen) and stained with the indicated antibodies in permeabilization buffer for 30 min at 4°C in the dark. For intracellular cytokine staining, cells were simulated ex vivo with PMA (Millipore-Sigma;10ng/ml) and ionomycin (Millipore-Sigma; 1 µM) for 4 h before staining, with monensin (Calbiochem; 2 µM) added during the final 2 h. Data were acquired on a FACSymphony A1 flow cytometer (BD Biosciences) and analyzed using FlowJo software.

### Co-immunoprecipitation

iTreg cells of the indicated genotypes or knockdown conditions were lysed in Pierce IP lysis buffer (Thermo Scientific) supplemented with PMSF protease inhibitor (Thermo Scientific). Lysates were clarified by centrifugation and equal amounts of protein (500 µg per sample) were incubated with anti-Lag3 antibody (R&D Systems; 5 µg) or an isotype-matched IgG control overnight at 4°C with rotation. Immune complexes were captured using protein A/G beads (Santa Cruz) for 2 h at 4°C with rotation. Beads were washed 3 times with Pierce IP lysis buffer, and bound proteins were eluted in Laemmli sample buffer. Immunoprecipitates were analyzed for CIN85 and HPK1, and input lysates were analyzed for Lag3 by western blotting as described below.

### Immunoblotting

Protein samples were denatured at 95°C for 10 min using 4x Laemmli buffer (Bio-Rad), resolved by SDS-PAGE, and transferred to nitrocellulose membranes (Whatman).

Membranes were blocked for 1 h and incubated with primary antibodies overnight at 4°C. Following washing, membranes were incubated with the corresponding secondary antibodies (Cell Signaling Technology) for 1 h at room temperature. Protein bands were detected using Pierce ECL Plus Western Blotting Substrate (Fisher Scientific) and visualized on iBright CL750 (Invitrogen).

### Quantitative RT-PCR

Total RNA was isolated using TRIzol reagent (Invitrogen) according to the manufacturer’s instructions and quantified using a NanoDrop spectrophotometer (Thermo Scientific). cDNA was generated using M-MLV reverse transcriptase (Promega). Quantitative real-time PCR was performed using Radiant qPCR Master Mix (Alkali Scientific) on a QuantStudio 3 Real-Time PCR System (Applied Biosystems).

Target gene expression was normalized to *Gapdh*, and relative expression was calculated using the 2⁻ΔΔCt method.

### Seahorse

Cells were analyzed for ECAR using a Seahorse XFe96 analyzer. Plates were pretreated with poly-D-lysine for cell adhesion, after which 2.5x10^5^ cells were plated per well. After adhesion and calibration, ECAR was measured using glucose (50 mM), oligomycin (1 mM), and 2-DG (200 mM).

### Nucleofection

Sorted naïve CD4^+^ T cells were washed with PBS, and 3x10^6^ cells per reaction were resuspended in 100 µL of electroporation buffer (Mirus Bio). Cas9 ribonucleoprotein (RNP) complexes were assembled by combining 2 µL of target-specific or scrambled control sgRNA (0.15 nmol/µL), 0.6 µL of Cas9 protein (IDT; 10 mg/ml), and 2.4 µL of nuclease-free water, followed by incubation for 10 min at room temperature. Cells were gently mixed with the RNP complexes, transferred to 0.2-cm electroporation cuvettes (Mirus Bio), and nucleofected using a Nucleofector II/2b (Lonza), with the mouse T cell program X-001. Immediately after nucleofection, 500 µL of prewarmed complete RPMI was added to each cuvette. Cells were transferred to 24-well plates containing 1.5 ml of prewarmed complete RPMI per well and allowed to recover for approximately 2 h at 37°C. Cells were then collected, washed with PBS, and cultured under iTreg-polarizing conditions for 3 days.

### Statistical analysis

Statistical analyses were performed using GraphPad Prism version 10 (GraphPad Software). Data are presented as mean ± SEM unless otherwise indicated. Statistical significance was determined by two-tailed unpaired t test with Welch’s correction, two-tailed Mann–Whitney U test, one-way ANOVA with Dunnett’s multiple-comparisons test, and two-way repeated-measures ANOVA with Šídák’s multiple comparisons test as indicated in the figure legends. P < 0.05 was considered statistically significant.

