## Supplementary Figures for "The Lag3 KIEELE motif couples HPK1 to metabolic and functional regulation of Foxp3+ regulatory T cells"

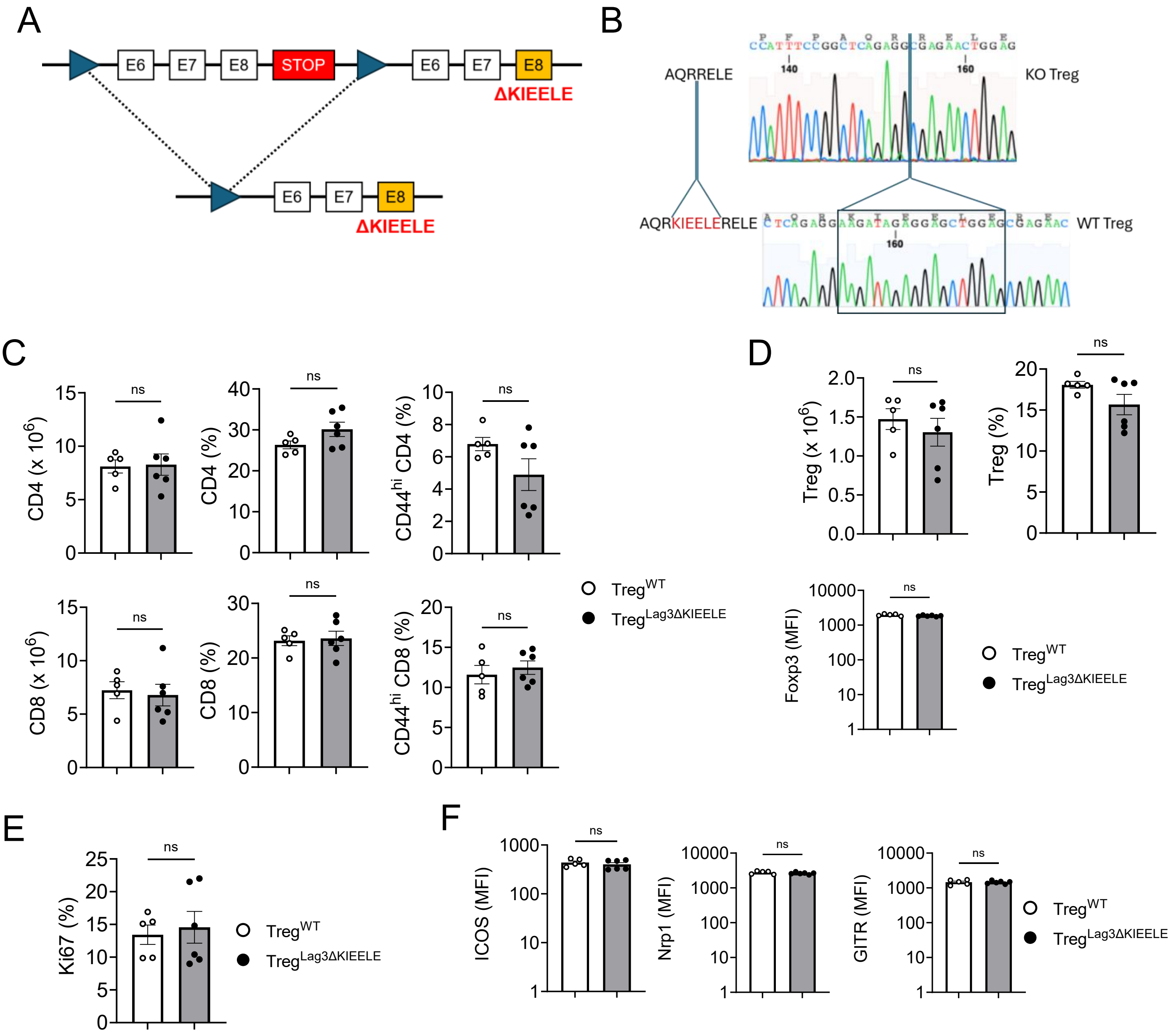


**Figure S1. Generation and steady-state characterization of mice with Treg-specific deletion of the Lag3 KIEELE motif.**

(A) Generation of Treg-specific Lag3 KIEELE mutant mice. (B) Genomic DNA sequencing of $Treg^{WT}$ and $Treg^{Lag3ΔKIEELE}$ mice. (C) Absolute number and proportion of CD4 and CD8 T cells and proportion of $CD44^{high}$ memory phenotype CD4 and CD8 T cells in the lymph node (LN). (D) Absolute number, proportion, and mean fluorescence intensity (MFI) of Foxp3 of $CD4^{+}$ $Foxp3^{+}$ Treg cells. (E) Proportion of $Ki67^{+}$ cells among $CD4^{+}$ $Foxp3^{+}$ Treg cells. (F) MFI of ICOS, Nrp1, and GITR in $CD4^{+}$ $Foxp3^{+}$ Treg cells in the LN. Statistical significance was determined using a two-tailed unpaired t test with Welch's correction. ns = not significant.

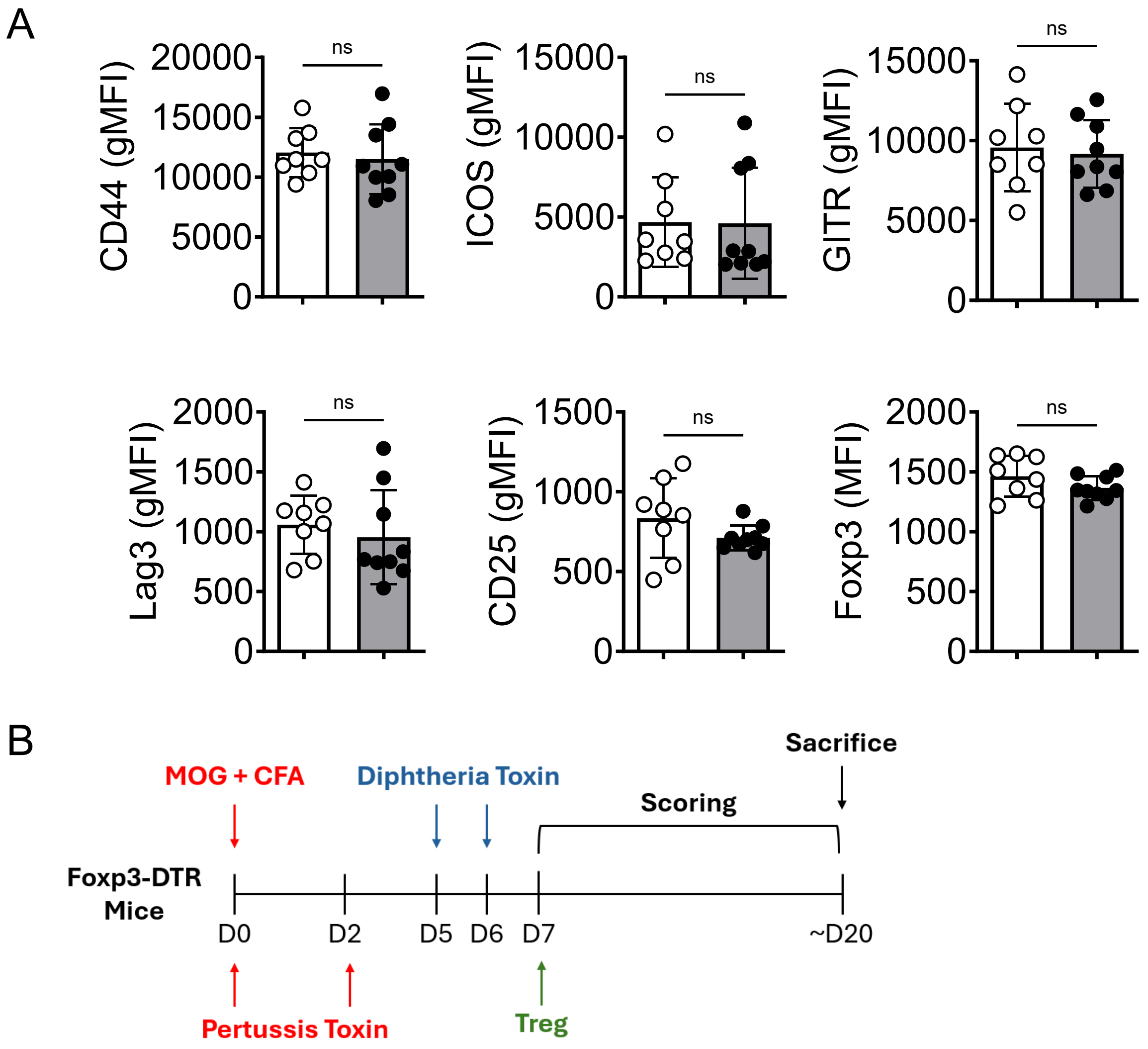


**Figure S2. CNS-infiltrating Treg characterization and experimental design for Treg adoptive transfer.**

(A) Geometric mean fluorescence intensity (gMFI) of CD44, ICOS, GITR, Lag3, CD25, and Foxp3 in CD4⁺ Foxp3⁺ Treg cells from the CNS at the peak of disease. (B) Experimental timeline of EAE induction, diphtheria toxin administration, and Treg adoptive transfer. Data were pooled from at least two independent experiments. Statistical significance in for experiments was assessed using a two-tailed Mann–Whitney U test. ns = not significant.

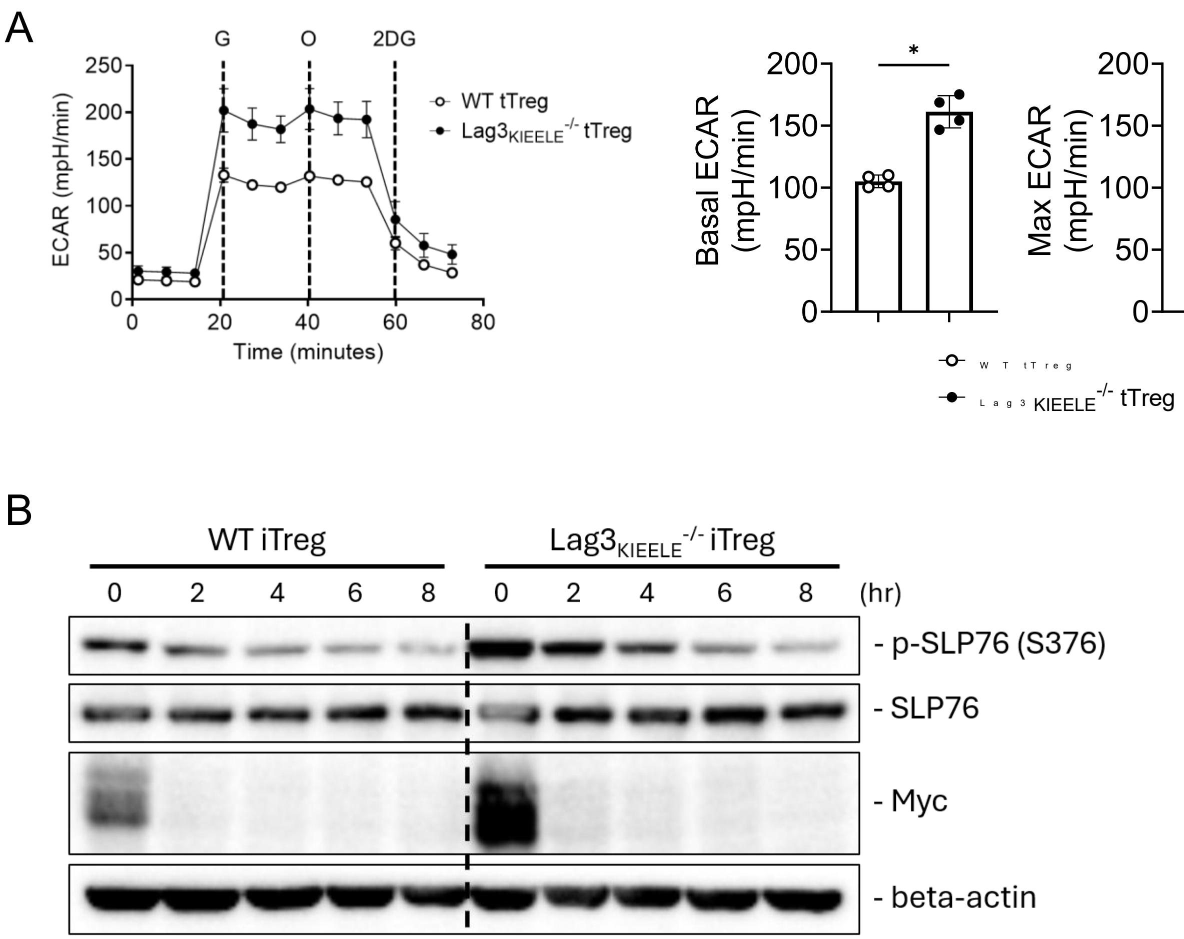


**Figure S3. Glycolytic activity in Lag3$_{KIEELE}$$^{-/-}$ tTreg cells and analysis of SLP76 phosphorylation and protein stability in iTreg cells.**

(A) ECAR of WT and Lag3$_{KIEELE}$$^{-/-}$ tTreg cells was measured by Seahorse. (B) WT and Lag3$_{KIEELE}$$^{-/-}$ iTreg cells were treated with cycloheximide for the indicated times (hr). Protein levels of p-SPL76 (S376), total SLP76, and Myc were assessed by western blotting. Data are representative of at least two independent experiments. Statistical significance was assessed using a two-tailed Mann–Whitney U test. *$P < 0.05$.

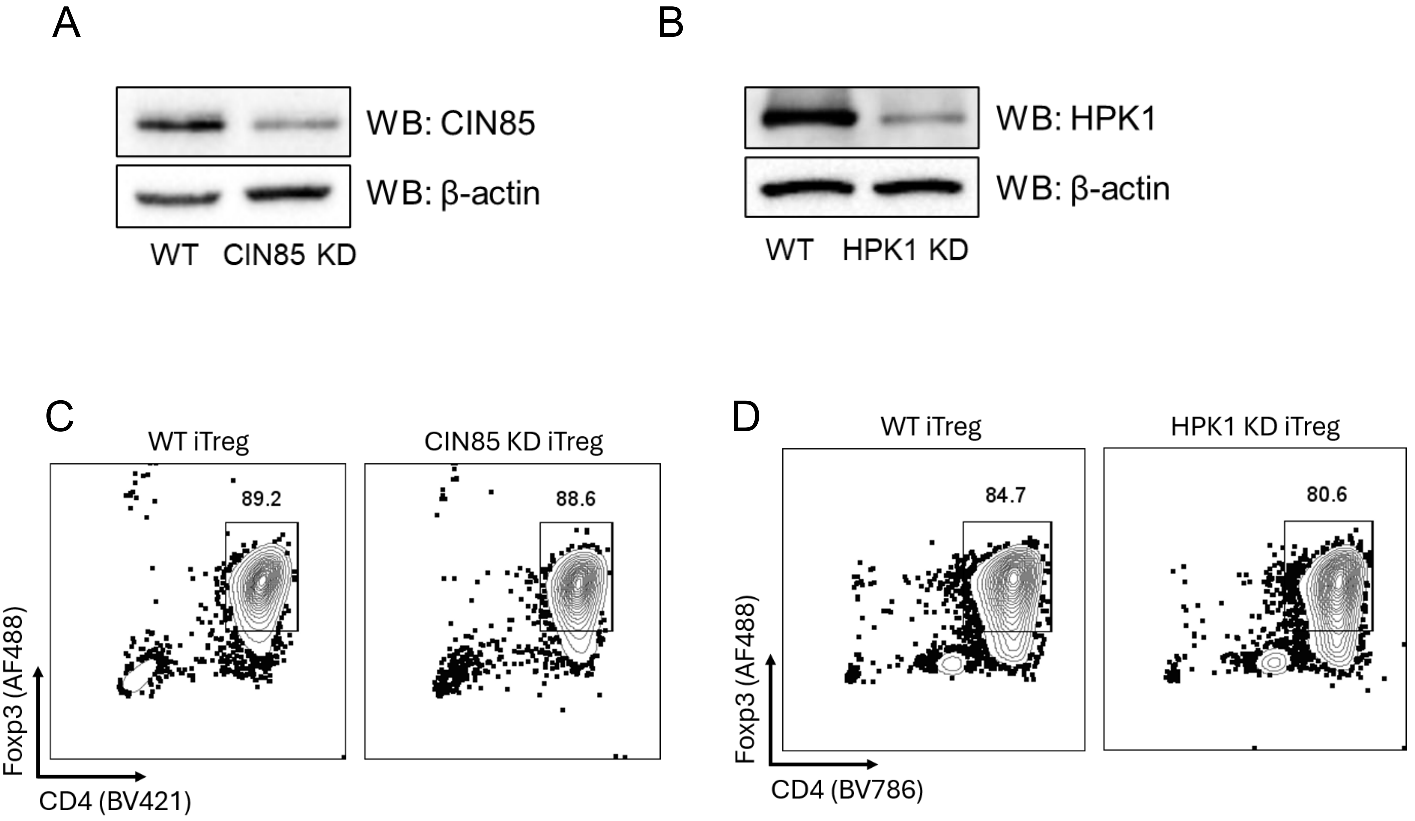


**Figure S4. Validation of CIN85 and HPK1 knockdown and assessment of Foxp3 expression in KD Treg cells.**
(A, B) WT naïve CD4 T cells were subjected to CRISPR-Cas9-mediated knockdown of CIN85 (A) or HPK1 (B) and cultured in Treg-polarizing condition for 3 days. CIN85 (A) or HPK1 (B) protein levels were assessed by western blotting. (C, D) WT and CIN85 KD iTreg cells (C) or WT and HPK1 KD (D) iTreg cells were stained for CD4 and Foxp3 and analyzed by flow cytometry to assess Foxp3 expression. Data are representative of at least two independent experiments.

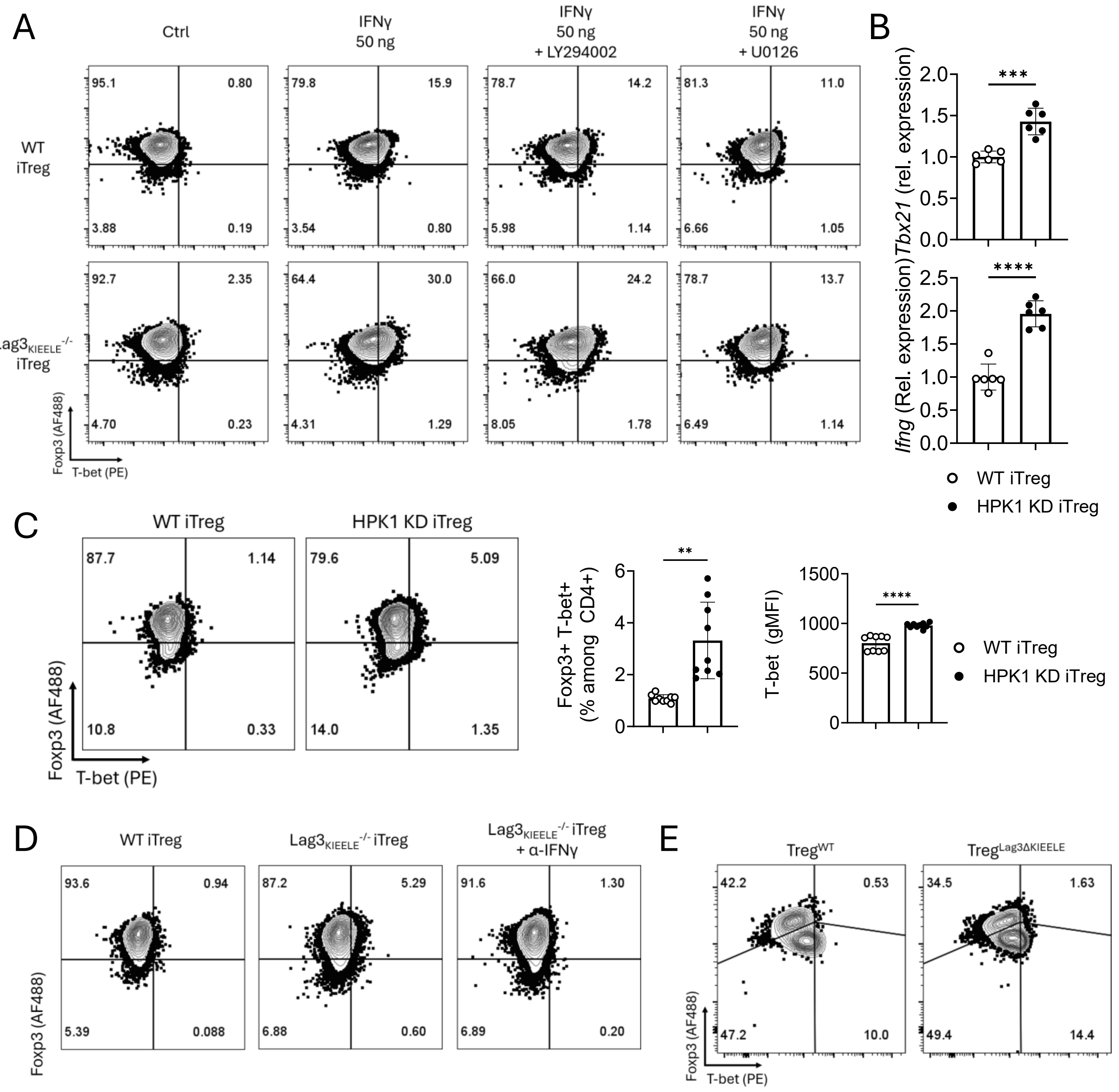


**Figure S5. Representative flow cytometry plots supporting Figure 4 and analysis of T-bet expression following HPK1 depletion in iTreg cells.**

(A, D, E) Representative flow cytometry plots corresponding to Figure 4B (A), Figure 4C (D), and Figures 4F and 4G (E). (B, C) mRNA expression of *Tbx21* and *Ifng* (B) and T-bet protein expression (C) measured from WT and HPK1 KD iTreg cells. Data from (A, B, D, E) are representative of at least two independent experiments and data from (C) were pooled from two independent experiments. Statistical significance was determined by two-tailed unpaired t test with Welch's correction. $^{**}P < 0.01$; $^{***}P < 0.001$; $^{****}P < 0.0001$.
